## Supplementary Figures for "Decorin, a novel negative modulator of E-cadherin in inflammatory breast cancer"

Extended Data [Supplementary] Figures 1-10

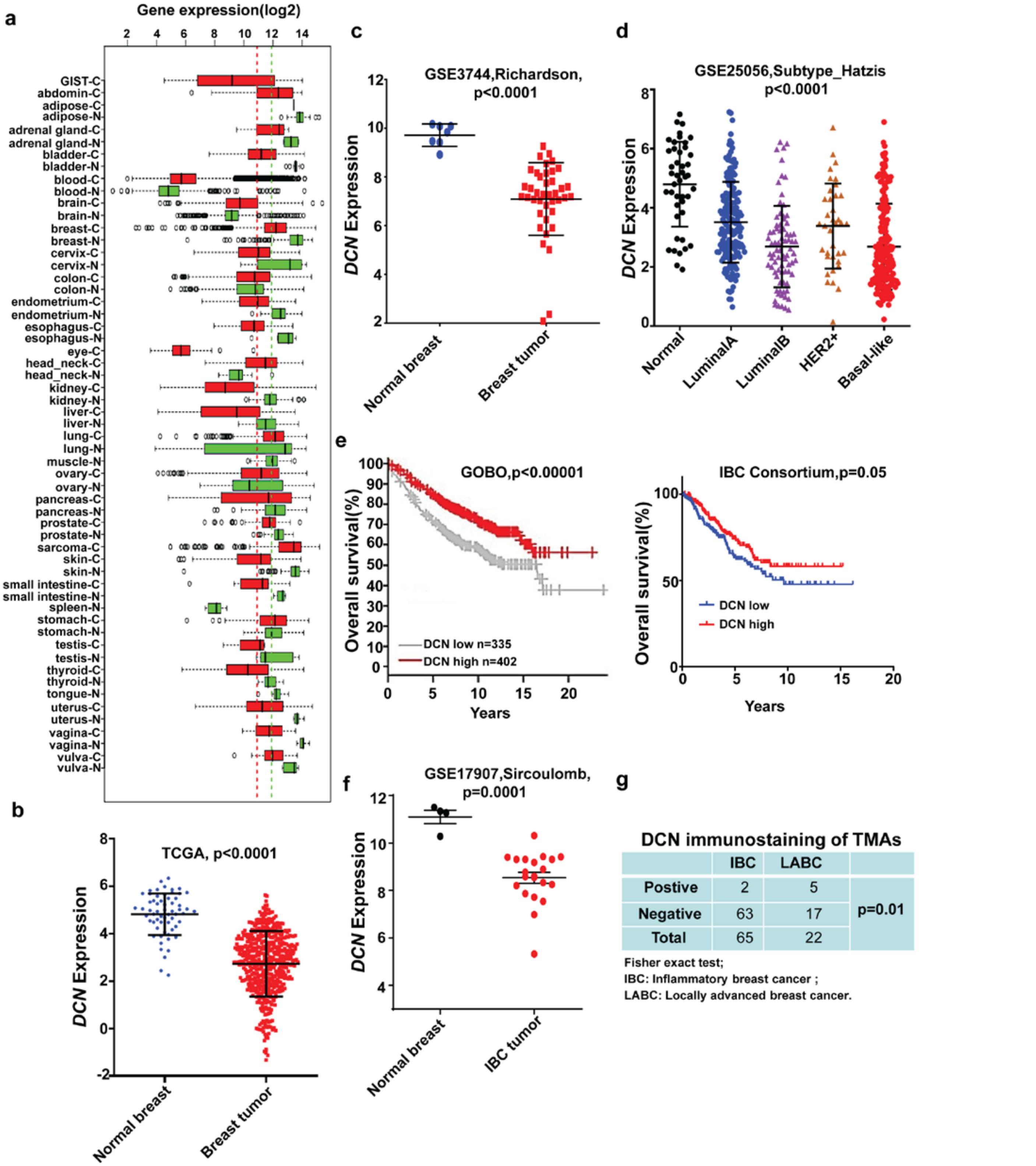


DCN immunostaining of TMAs

| | IBC | LABC | |
|---|---|---|---|
| Postive | 2 | 5 | p=0.01 |
| Negative | 63 | 17 | |
| Total | 65 | 22 | |

Fisher exact test;
IBC: Inflammatory breast cancer ;
LABC: Locally advanced breast cancer.

**[Supplementary] Extended Data Figure 1: DCN is downregulated in aggressive breast tumors.**

**a,** Decorin (*DCN*) mRNA expression pattern was analyzed in normal and tumor tissues across cancer types from the online database Gene Expression across Normal and Tumor tissues (GENT), which contains more than 34,000 samples. N, normal; C, cancer. **b and c,** DCN is downregulated in breast tumors relative to normal tissues, as shown in The Cancer Genome Atlas (TCGA) **(b)** and Richardson dataset (GSE3744) **(c)**. **d,** DCN expression is downregulated in more aggressive, basal-like breast cancer subtypes (Hatzis dataset, GSE25066). **e,** High DCN expression is associated with better survival outcomes, as indicated by the Gene Expression Based Outcome (GOBO) and Inflammatory Breast Cancer (IBC) Consortium datasets. **f,** DCN expression is downregulated in IBC compared with normal breast tissues, as indicated by the Sircoulomb dataset (GSE17907). **g,** DCN is expressed less frequently in the more aggressive IBC tumors than in locally advanced non-IBC breast cancer (LABC).

a

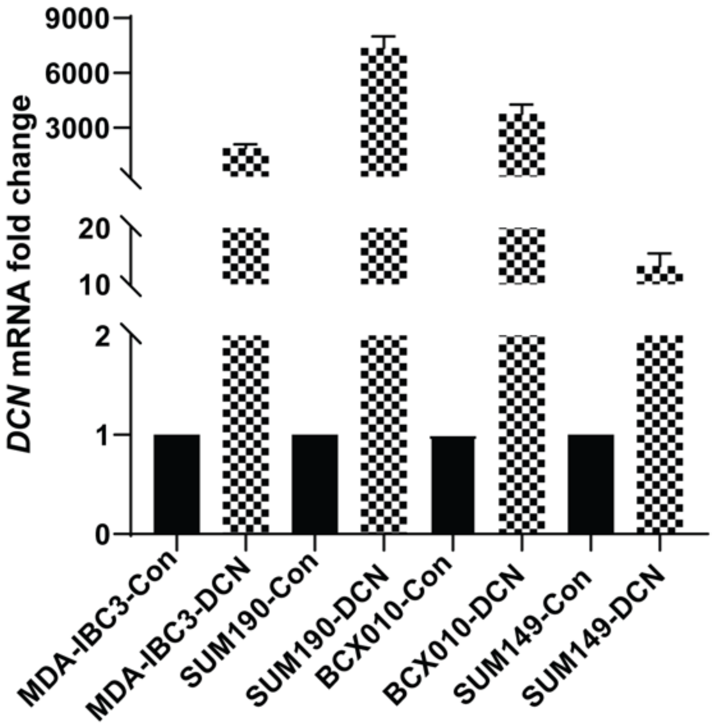


**[Supplementary] Extended Data Figure 2: Validation of DCN mRNA overexpression in IBC cell lines.** *DCN* mRNA levels were assessed by quantitative RT-PCR in DCN-overexpressing and control IBC cell lines. Data are from three independent assays.

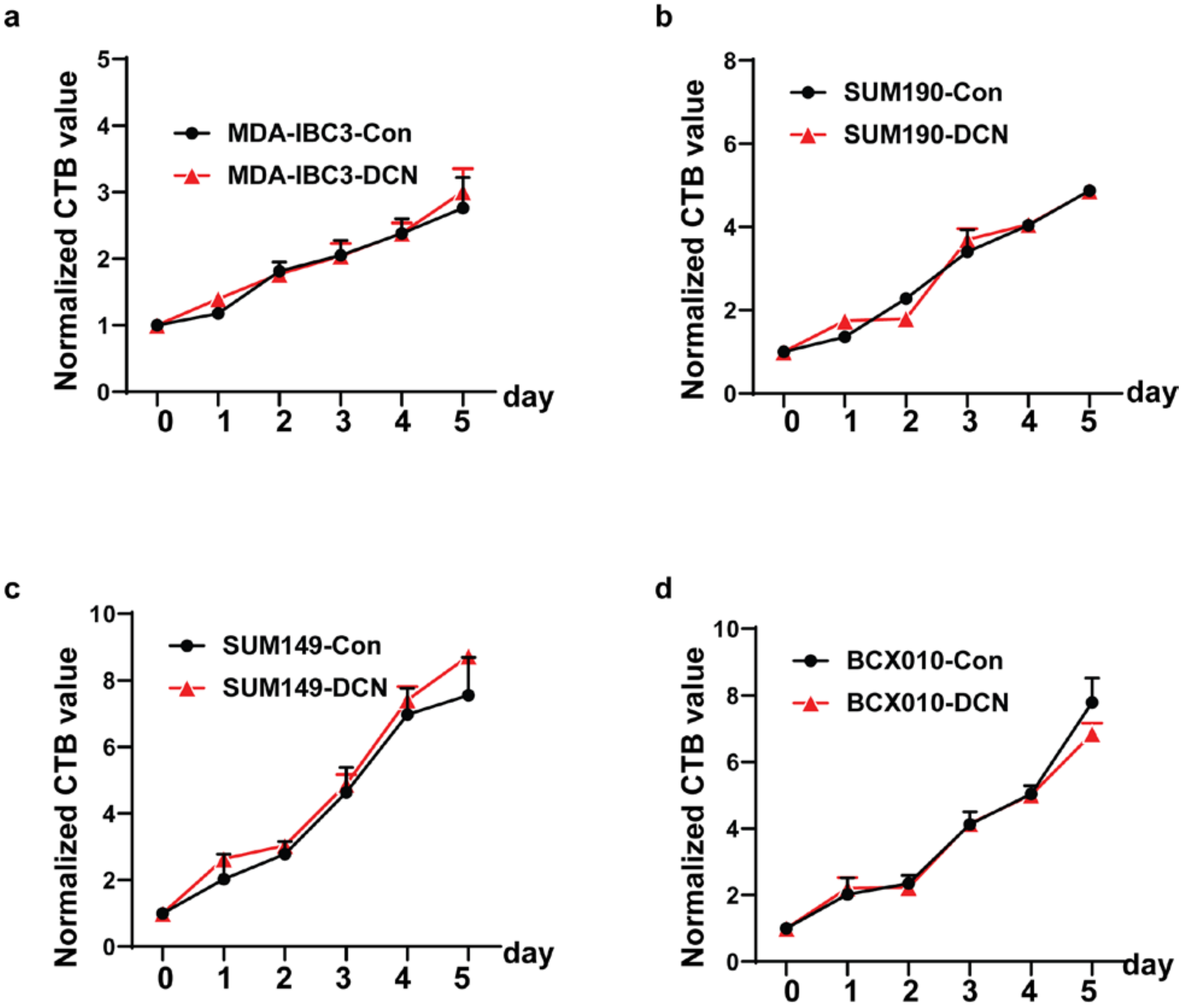


**[Supplementary] Extended Data Figure 3: DCN does not affect proliferation in IBC cell lines.** Cell number was evaluated with a CellTiterBlue assay (Promega) on the indicated days. The fluorescence value of the cells measured at the start of the experiment was set as 1. Each cell line was evaluated separately at the indicated periods, and the fluorescence values of treated cells were normalized to their corresponding non-treated controls. **a-d,** Results in MDA-IBC3 cells (**a**), SUM149 cells (**b**), SUM190 cells (**c**), and BCX010 cells (**d**) show no significant differences between the DCN-overexpressing and control groups. Results were normalized to the controls. All data are represented as means ± standard error of the mean, with all experiments done in triplicate.

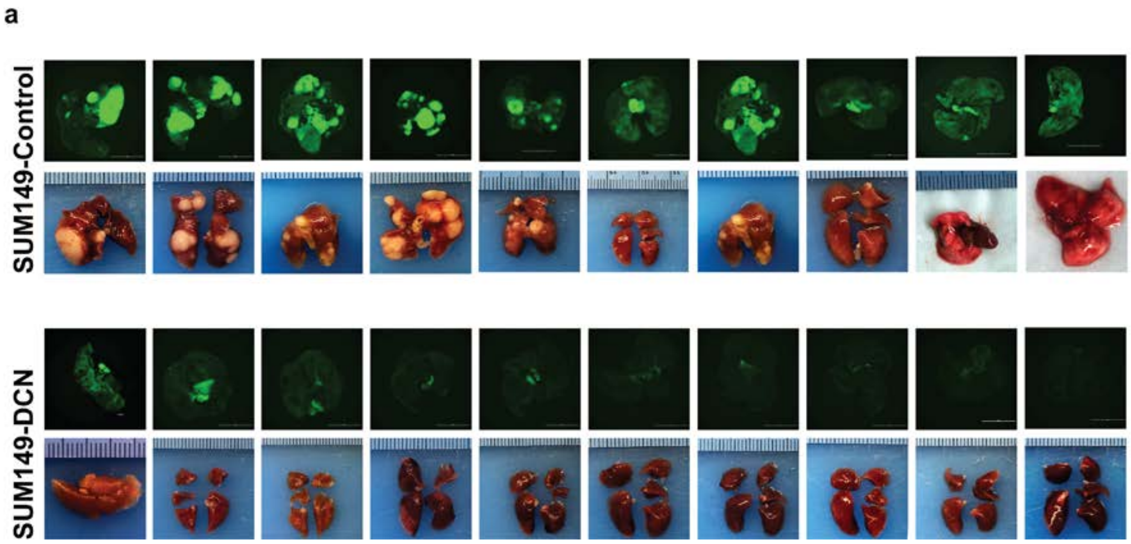


**[Supplementary] Extended Data Figure 4: DCN overexpression inhibits lung metastatic colonization *in vivo*** GFP-labeled DCN-overexpressing or control SUM149 cells were injected via tail vein into SCID/Beige mice (10 mice/group). Images of lungs from all of the mice in each group are shown here.

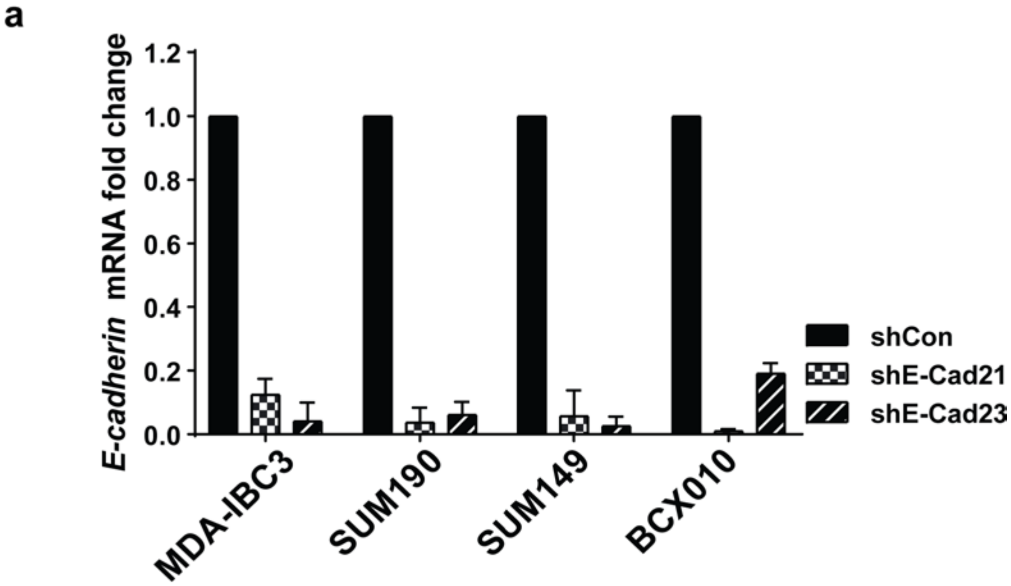


**[Supplementary] Extended Data Figure 5: Validation of E-cadherin mRNA knockdown in IBC cell lines.** *E-cadherin* mRNA levels were measured by quantitative RT-PCR in four cell lines with E-cadherin knockdown. Data are from three independent assays.

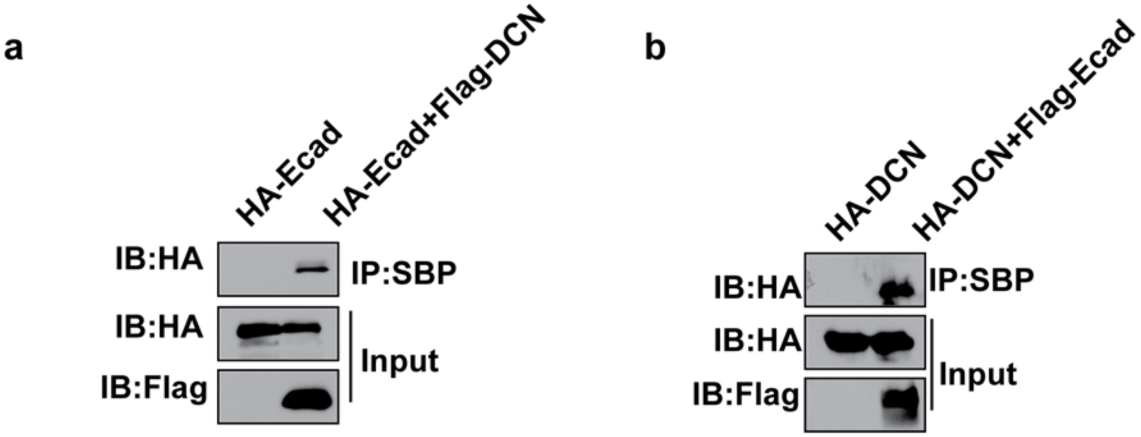


**[Supplementary] Extended Data Figure 6: DCN forms a complex with E-cadherin in HEK293T cells *in vitro*. a,** HA-E-cadherin and Strep-E-cadherin-flag-DCN were co-transfected into 293T cells. **b,** HA-DCN and HA-DCN-flag-E-cadherin were co-transfected into HEK293T cells. Cell lysates were analyzed by immunoprecipitation and western blotting with anti-FLAG and anti-HA antibodies. Whole-cell lysates were blotted and shown as the input.

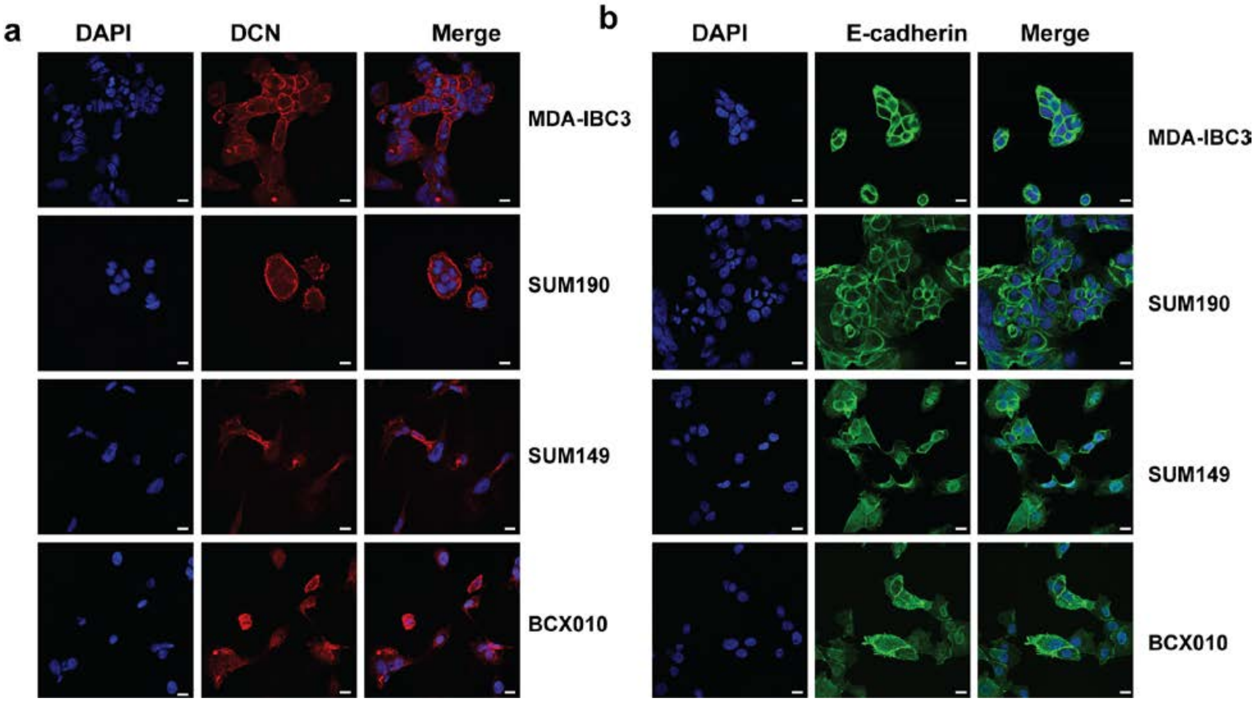


**[Supplementary] Extended Data Figure 7: DCN and E-cadherin are co-localized on the membrane.** Immunofluorescence staining of endogenous DCN and E-cadherin in IBC cell lines. **a and b,** Single-immunofluorescence staining shows localization of DCN (red) (**a**) and E-cadherin (green) (**b**) on the membranes of IBC cells.

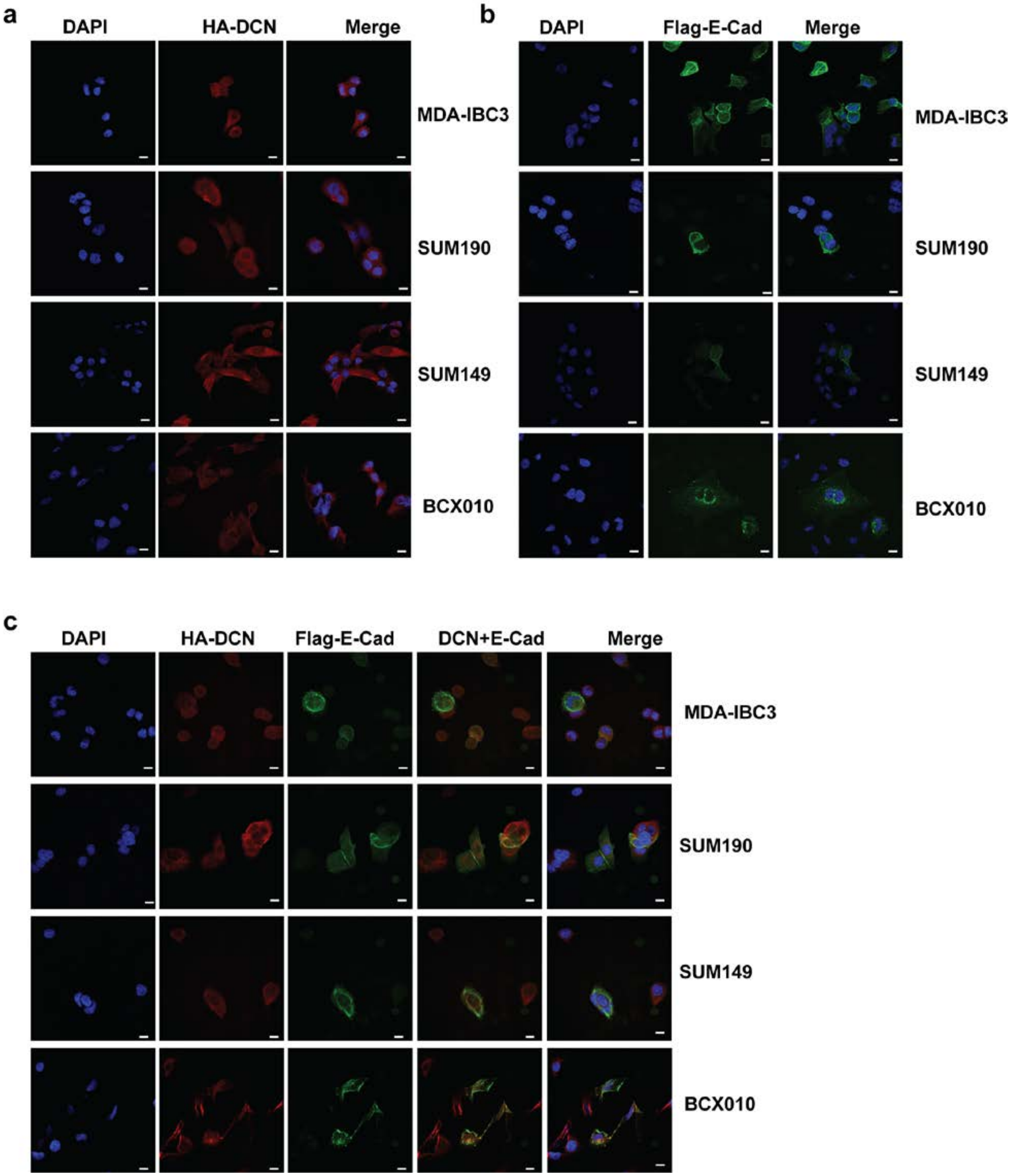


**[Supplementary] Extended Data Figure 8: HA-Decorin and Strep-flag-E-cadherin are co-localized on cell membranes. a and b**, HA-DCN or strep-flag-E-cadherin was transfected individually into each of the indicated IBC cell lines, and subcellular localization was examined by using anti-HA polyclonal antibody (red) or anti-Flag monoclonal antibody (green). Both were localized on the membranes of IBC cells. **c,** HA-DCN and Strep-flag-E-cadherin were co-transfected into IBC cells and the subcellular localization of both flags was examined by using an anti-HA polyclonal antibody (red) and anti-Flag monoclonal antibody (green). Both HA-DCN and Flag-E-cadherin were co-localized on the cell membranes.

**a**

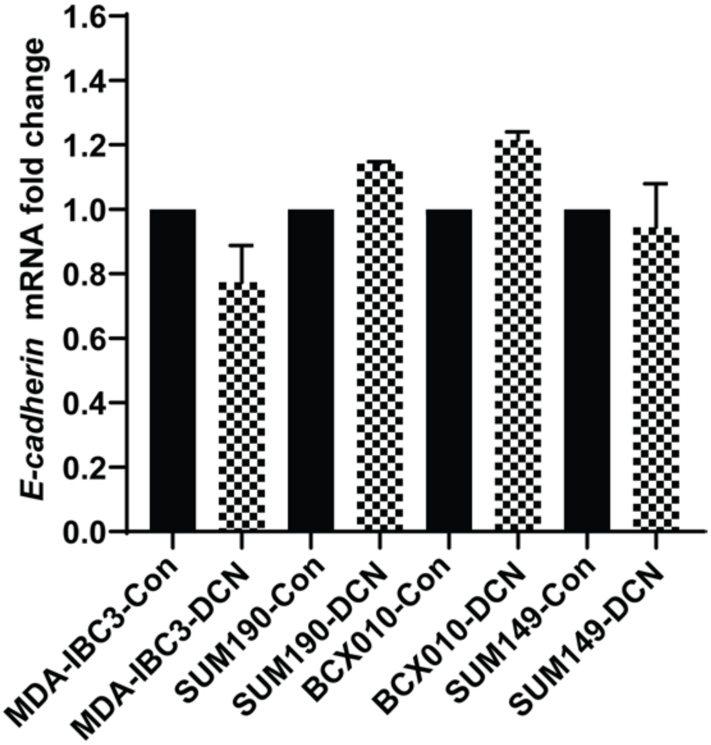


**[Supplementary] Extended Data Figure 9: DCN overexpression does not affect *E-cadherin* mRNA expression.** *E-cadherin* mRNA levels were quantified by qRT-PCR in control and DCN-overexpressing IBC cell lines.

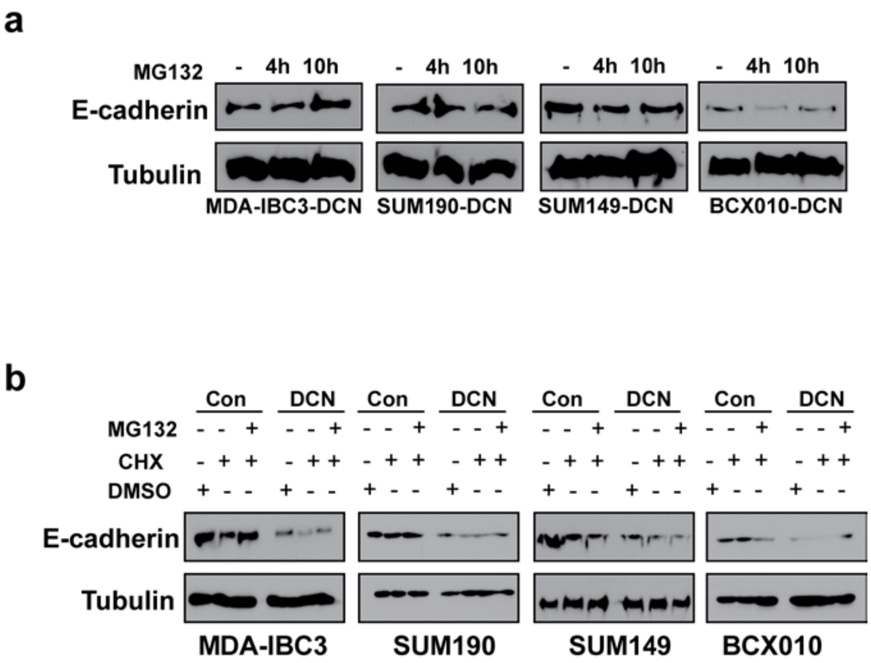


**[Supplementary] Extended Figure 10: Proteasome inhibition by MG132 treatment does not rescue E-cadherin expression in DCN-overexpressing IBC cells.** Inhibiting proteasomes with the proteasome inhibitor MG132 does not block DCN-induced degradation of E-cadherin. **a,** DCN-overexpressing IBC cells treated with MG132 (50nmol) for 4 hours or 10 hours showed no differences in E-cadherin protein levels over time. **b,** DCN-overexpressing and control IBC cells were treated with the protein synthesis inhibitor cycloheximide (CHX,100μg/ml) with or without MG132 (50 nmol) for 10 hours. No differences in E-cadherin protein levels were noted between DCN-overexpressing and control cells. Collectively, these results indicate that DCN-mediated E-cadherin degradation is independent of proteasome activity.
